# Exploiting expression patterns across multiple gene isoforms to identify radiation response biomarkers in early-stage breast cancer patients

**DOI:** 10.1101/086322

**Authors:** Chaitanya R. Acharya, Kouros Owzar, Janet K. Horton, Andrew S. Allen

## Abstract

In an effort to understand the underlying biology of radiation response along with whole transcriptome effects of preoperative radiotherapy in early-stage breast tumors, we propose two efficient score-based statistical methods that exploit gene expression patterns across all available gene transcript isoforms and identify potential biomarkers in the form of differentially expressed genes and differentially enriched gene-sets. We demonstrate the effectiveness of these two methods using extensive simulation studies that show that both of our methods give improved performance, in terms of statistical power, over the most commonly used methods. By exploiting radiation-induced changes in all available gene transcript isoforms, we identified several statistically significant differentially expressed genes related to PI3K-AKT and JAK-STAT signaling pathways along with radiation-induced oncogenic signaling pathways and tumor microenvironment gene signatures that could be potential targets to improve response to radiotherapy in breast tumors.

## Background

Radiation therapy or radiotherapy is utilized as a curative therapy in many solid tumors including gynecologic, head and neck, gastrointestinal, breast, prostate, lung, central nervous system and pediatric malignancies. Approximately 60% of cancer patients receive radiotherapy as part of their treatment either as a stand alone pre-operative therapy or combined with other modalities such as chemotherapy following surgery in an adjuvant setting [1]. Radiotherapy has played a significant role in treating both invasive and non-invasive breast tumors over the years. However, response to radiation in breast cancer patients has not been uniform across all breast tumor subtypes (for example, basal, luminal, etc.) leading to a significant percentage of patients being either over- or under-treated [2, 3]. This can be attributed to variable transcriptional response (through variable activation of transcription factors) to radiation, which is very similar to response to chemotherapy except that the mechanisms underlying radiation response have not been well understood and studied [4, 5]. Many genes have multiple transcript isoforms that result from alternative-splicing events. We measure the overall gene expression of a given gene by measuring the relative abundances of these isoforms, which may provide new insights into disease and biology.

Constantly evolving high-throughput gene expression profiling technologies, such as RNA-Seq or ultra high-resolution microarrays, have enabled us to interrogate all transcript isoforms in the human transcriptome by targeting coding transcripts, exon-exon splice junctions, and non-coding transcripts. The end goal of using these technologies is to exploit the gene expression patterns across multiple isoforms or gene transcripts in order to map biomarkers such as genes and gene-sets that help illuminate the molecular pathology of complex diseases at the RNA level. Existing analytic tools or methods for biomarker analysis such as differential expression analysis involves combining gene expression over all gene isoforms prior to data analysis, resulting in a gene-level interrogation of biological conditions [6, 7, 8, 9, 10, 11]. For example, the overall expression level of a gene can be represented by a single number and is measured by averaging the signals of many transcripts for the gene. Individual transcripts that have high variability compared to the average expression of a gene will be removed from the analysis (outliers). Such an approach has at least two significant limitations. First, it fails to fully exploit expression patterns across gene isoforms either by combining information across multiple transcripts or by not explicitly identifying effects that differ across transcripts. Second, and more importantly, it fails to account for alternative splicing or alternative 3′ poly-adenylation events by removing gene isoforms that seems to be significantly differentially expressed. We propose two distinct approaches, one to identify radiation-induced gene expression biomarkers in an isoform-specific differential expression (DE) analysis and another to perform isoform-specific gene-set or pathway enrichment analysis. We test these methods extensively using simulation studies and then evaluate the effectiveness of these two methods on a microarray-based gene expression dataset containing 26 paired early-stage breast cancer patient samples. Briefly, these tumor samples originated from a unique preoperative radiotherapy Phase I trial [2], and were assayed on the new Affymetrix Human Transcriptome 2.0 array [29]. The transcriptome response to radiation exposure was derived by comparing gene expression in samples before and after irradiation. While demonstrating the effectiveness of our method to identify differentially enriched gene-sets, we investigated the effects of radiation on 7 tumor microenvironment and 24 hallmark oncogenic signaling gene-sets that are associated with radiation response.

We hypothesize that our methods are effective in identifying biomarkers (in this case, differentially expressed genes and differentially enriched gene-sets) when compared to most commonly used approaches. Investigating the tumor microenvironment and the oncogenic signaling pathways before and after radiation will help us understand any radiation-induced changes in individual patients, which may serve as a surrogate to understand patient response to radiation and can make for potential therapeutic targets.

## Materials and Methods

### Microarray analysis of the breast cancer dataset

Raw microarray data for twenty six early-stage breast cancer patients were obtained from NCBI’s gene expression omnibus (GEO ID: GSE65505) repository [28]. All the patients are at least 55 years old, clinically node negative, ER-positive and/or PR-positive, HER2-negative (biologically favorable tumors) with T1 invasive carcinomas or low-intermediate grade *in situ* disease ≤ 2*cm*. These patients received pre-operative radiotherapy (radiation dose prior to surgical resection of tumor). All the samples were arrayed on Affymetrix Human Transcriptome Array 2.0 [29], which was designed with approximately ten probes per exon and four probes per exon-exon splice junction. At the top level, each transcript cluster roughly corresponds to a gene. Each transcript cluster is comprised of exon clusters that a) shared splice sites, b) or were derived from overlapping exonic sequences, c) or were single-exon clusters bounded on the genome by spliced content. Each exon cluster is further fragmented into probe selection regions (PSRs), which are non-overlapping contiguous sequences. Gene-level and gene isoform/transcript-level expression data were obtained using R/Bioconductor packages *oligo* [30], *affyio* [31] and *pd.hta.2.0* [32], and pre-processed by robust multi-array average (RMA) method [7, 30, 31], which summarizes the probe level expression data into a probe set level expression value. Principal component analysis was conducted to check for batch effects in both gene-level and transcript-level data, and any batch effects that were identified were corrected using a popular Empirical Bayes approach (ComBat) [33]. DE analysis was performed on genes with at least two transcript isoforms. This resulted in a dataset with more than 800,000 transcripts.

### Strategy to identify gene expression biomarkers of radiation: Differential Expression (DE) analysis

Given two distinct biological groups (before and after radiation treatment), gene expression for each gene transcript, *Y*, can be modeled in the following way

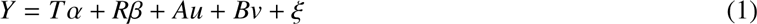

where *Y* is a *ntg* × 1 matrix of expression values, *T* is a *ntg* × *t* dimensional matrix of gene expresion levels in *t* isoforms of a gene in *g* groups and *n* individuals, α is a fixed effect representing the isoform-specific intercepts, *R* is a *ngt* × 1-dimensional matrix of radiation dose identifiers such that *R* ∈ {0, 1}, 0 indicates no radiation and 1 indicates radiation, β is a fixed effect indicating the average effect of radiation on gene expression. *u* ∼ *N* (0, τ*AA^T^*) indicates subject-specific random intercept, *ν* ∼ *N* (0, γ*BB^T^*) is random effect that denotes the interaction between gene-isoform and radiation (isoform-specific radiation effect), and *ξ* ∼ *N*(0, *∈I*). *I* is *ntg* × *ntg* dimensional identity matrix. The matrices *J*, *A* and *B* are design matrices with *B* being a function of radiation dose. *J* is *ntg* × *t* dimensional matrix denoting the design matrix for the tissue-specific intercepts. *A* is *ntg* × *n* design matrix for the subject-specific intercepts. *B* is a *ntg* × *t* design matrix of stacked radiation dose identifiers.

We test the null hypothesis that *H*_0_ : *β* = 0; γ = 0 i.e radiation does not affect gene expression. From our model above, we derive our score test statistic, *U*_*ψ*_ as

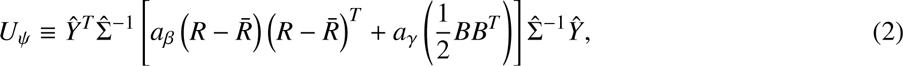

where *a*_*β*_ and *a*_*γ*_ are scalar constants chosen to minimize the variance of *U*_*ψ*_ (see Supplementary methods). The *p* values are approximated using Satterthwaite method [34]. The maximum likelihood estimates, obtained from fitting a standard linear mixed model using lme4 [35], are computed only once per gene since under the null, there is no effect due to radiation on the gene expression. The *p* values obtained from applying our method were adjusted for multiple hypothesis within the false discovery rate (FDR) framework. Genes with FDR adjusted *p* values (*q* values) less than 0.05 were selected to be differentially expressed. More information on our method is available in the supplementary methods. As a side note, this model is very similar to a previous one we proposed [36] with the exception that this is a paired data.

### Strategy to perform radiation-induced isoform-specific gene-set enrichment analysis

Gene expression data for each pathway, *Y*, is modeled in the following way

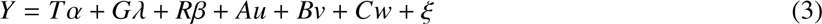

where *Y* is *nt jg* × 1 dimensional matrix of expression values, *T* is a *nt jg* × *t*-dimensional matrix of expression levels in *t* isoforms of a gene, *j* genes, *g* groups and *n* individuals, α is a fixed effect representing *t* isoform-specific intercepts, λ is a fixed effect representing *g* gene-specific intercepts, *R* is a *nt jg* × 1 dimensional matrix of radiation dose identifiers such that *R* ∈ {0, 1}, 0 indicates no radiation and 1 indicates radiation, *β* is a fixed effect indicating the average effect of radiation on a pathway or gene-set. *υ* ∼ *N* (0, τ*AA_T_*) indicates subject-specific random intercept, *w* ∼ *N* (0, γ*BB_T_*) is a random effect that denotes the interaction between gene-isoform and radiation (isoform-specific radiation effect), *ω* ∼ *N* (0, φ*CC_T_*) is a random effect that denotes the interaction between gene and radiation (gene-specific radiation effect), and *ξ* ∼ *N* (0, *∈I*). *I* is *nt jg* × *nt jg*-dimensional identity matrix ∙ The matrices *J*, *A* and *B* are design matrices with *B* being a function of radiation dose. *J* is *nt jg* × *t* dimensional matrix denoting the design matrix for the tissue-specific intercepts. *A* is *nt jg* × *n* design matrix for the subject-specific intercepts. *B* is a *nt jg* × *t* design matrix of stacked radiation dose identifiers and *C* is a *nt jg* × *g* dimensional design matrix of the *R* × *G* effect.

We test the null hypothesis that *H*_0_ : *β* = 0; γ = 0; *φ* = 0 i.e radiation does not affect gene expression. From our model above, we derive our score test statistic, *U_ζ_* as

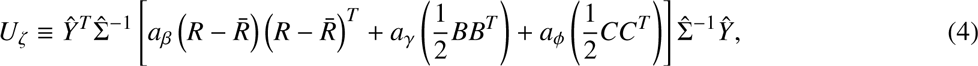

where *a_β_*, *a_γ_* and *a_φ_* are scalar constants chosen to minimize the variance of *U_ζ_*. The *p* values are approximated using Satterthwaite method [34]. Similar to our earlier method, the maximum likelihood estimates, obtained from fitting a standard linear mixed model using lme4 [35], are computed only once per gene-set since under the null, there is no effect due to radiation on the gene expression. The *p* values obtained from applying our method were adjusted for multiple hypothesis within the false discovery rate (FDR) framework. Genes with FDR adjusted *p* values (*q* values) less than 0.05 were selected to be differentially expressed. More details on our method are available in supplementary methods.

## Simulations

### Testing our method for DE analysis

We have performed the following two simulation studies in order to verify our approach. In our first study, we simulated one gene at a time from the following linear model and varied the following parameters-*β* (additive effect due to radiation), the proportion of variation explained by γ or *R* ×*T* effect 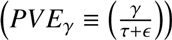 and the number of transcripts. For a positive integer *tg* that represents the combined number of transcripts (*t*) and groups (*g*), if **1** denotes a column vector of *tg* ones and 𝕀 denotes the corresponding *tg* × *tg* diagonal matrix, following the *tg*-variate normal law denoted by *N_tg_* [*µ*, Σ] with mean *µ* ∈ ℝ^*tg*^ and variance Σ ∈ ℝ*^tg×tg^*, expression levels of a target gene *j* by using the following vectorized form of the linear mixed model –

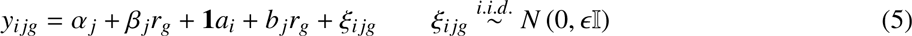

where *y_ijg_* is a *tg* × 1 vector of gene expression data, α*_t_* is the transcript-specific intercept (α*_t_* ∈ ℝ^*t*^), *β_j_* describes the main additive effect (*β_j_* ∈ ℝ^1^), *r*_*g*_ is a vector of length *tg* such that *r* ∈ (0_*t*_, 1_*t*_). The random effect *b_j_* ∈ ℝ^*tg*^ represents transcript-specific interaction effect of radiation, and *a*_*i*_ ∈ ℝ^1^ is a subject-specific random intercept. We assume that all the random effects are independent and that *a_i_* ∼ *N*_1_ (0, *τ*), *b_j_* ∼ *N_tg_* (0, *γ*𝕀). A linear mixed effects model was fit using the package *lme4* [35] in the statistical environment R (R Core Team).

We then compared our method with a standard paired t-test and a non-parametric alternative in Wilcoxon’s test [37]. The test statistic in case of transcript-by-transcript (TBT) analysis is the minimum *p* value over the total number of transcripts from either t-test or Wilcoxon’s test performed separately in each transcript for each paired sample. A gene-level test was constructed over all the transcripts by taking the median expression value across the transcripts followed by a standard paired t-test. Statistical significance was determined at a nominal *p* value of 0.05 for all power simulations (in case of TBT analysis, it is 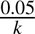 where *k* is the number of transcripts). We used 10,000 data replicates to evaluate the type I error and 1,000 data replicates for power calculations.

We have also tested our method on a synthetic dataset simulated from a multivariate normal distribution containing two classes of data. Each gene was simulated to have variable number of transcripts. We used this dataset with increasing number of genes (by also keeping a small proportion differentially expressed) and tested our approach at both transcript-level (paired t-test and Wilcoxon’s test) and gene-level. The most commonly used method to combine *p* values of all the transcripts of a gene is Fisher’s method however, under the assumption that all the *p* values are independent [14]. This assumption may be frequently violated since different isoforms of a gene may be correlated and the resulting *p* values are dependent on each other. At the gene-level, paired t-tests were run on gene expression values of a gene that were aggregated over its transcripts by either their median expression values or Winsorized mean [38] expression values.

### Testing our method for gene-set enrichment analysis

Similar to the above analyses, we have performed two simulations studies in order to verify our approach. In our first study, we simulated one gene-set at a time from the following linear model and varied the following parameters- *β* (additive effect due to radiation), the proportion of variation explained by *γ* or *R* × *T* effect 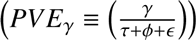 the proportion of variation explained by φ or *R* × *G* effect 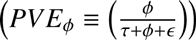 and the number of transcripts. For a positive integer *t jg* that represents the combined number of transcripts (*t*), genes ( *j*) and groups (*g*), if **1** denotes a column vector of *t jg* ones and 𝕀 denotes the corresponding *t jg* × *t jg* diagonal matrix, following the *t jg*-variate normal law denoted by *N_t jg_* [*µ*, Σ] with mean *µ* ∈ ℝ*^t jg^* and variance Σ ∈ R*t jg* ×*t jg*, expression levels of a target geneset *k* by using the following vectorized form of the linear mixed model –

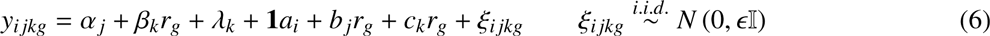

where *y_ijkg_* is a *t jg* ×1 vector of gene expression data, α*t* is the transcript-specific intercept (α*t* ∈ R*t*), β*k* describes the main additive effect (β*k* ∈ R1), *rg* is a vector of length *t jg* such that *r* ∈ (0*tg*, 1*tg*). The random effect *bt* ∈ R*t jg* represents transcript-specific interaction effect of radiation, the random effect *c j* ∈ R*t jg* represents transcript-specific interaction effect of radiation, and *ai* ∈ R1 is a subject-specific random intercept. We assume that all the random effects are independent and that *ai* ∼ *N* 1 (0, τ), *bt* ∼ *Ntg* (0, γI) and *c j* ∼ *N jg* (0, γI). A linear mixed effects model was fit using the package *lme4* [35] in the statistical environment R (R Core Team).

We then compared our method with a standard paired t-test and a non-parametric alternative in Wilcoxon’s test [37]. The test statistic in case of transcript-by-transcript (TBT) within a gene analysis is the minimum *p* value over the total number of transcripts and genes from either t-test or Wilcoxon’s test performed separately in each transcript for each paired sample. A gene-level test was constructed over all the transcripts by taking the median expression value across the transcripts followed by a standard paired t-test. Statistical significance was determined at a nominal *p* value of 0.05 for all power simulations (in case of TBT analysis, it is 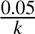, where *k* is the product of the number of transcripts and genes). We used 10,000 data replicates to evaluate the type I error and 1,000 data replicates for power calculations.

A second set of simulations involved generating a synthetic gene expression data from a multivariate normal distribution containing two classes of data. Each gene was simulated to have variable number of transcripts. We defined two types of gene-sets, one with overlapping genes and the other with non-overlapping genes, and randomly assigned some gene-sets to to contain differentially expressed genes. Since most, if not all of the current methods involve gene-set analysis at the gene level, we compared our method with Gene Set Variational Analysis (GSVA) [10], Pathway Level Analysis of Gene Expression (PLAGE) [39], single sample GSEA (ssGSEA) [40] and the combined z-score (ZSCORE) [41] methods. Both, PLAGE and the ZSCORE are parametric and assume that gene expression profiles are jointly normally distributed. More about these methods in the supplementary material.

### Defining the gene-sets and gene-set analysis

All the hallmark oncogenic signaling pathways used in our primary data analysis were obtained from the Molecular Signature Database version 3 (MSigDB) collection [20]. We focussed our attention on 24 specific oncogenic signaling pathways that were most likely associated with radiation response. We defined tumor microenvironment as a collection of proteins produced by cells present in and around the tumor that support the growth of the cancer cells. We included gene-sets representing hypoxia [42], invasiveness/metastases gene signature [43], epigenetic stem cell signature in cancer [44], inflammatory pathway involving tumor necrosis factors [45], angiogenesis [46], immune signatures [47] and a form of genomic instability called chromosomal instability [48], which determines the tumor cell’s ability to respond to its microenvironment. In order to visualize sample set enrichment of these gene-sets (enrichment level of a gene-set in a sample), we employed Gene Set Analysis (GSA) software [49], which implements a supervised method (class labels are known before the analysis) that computes a ”maxmean” summary statistic for each gene-set. Briefly, GSA computes the average of both positive and negative aspects of gene-scores (for example, fold changes) over each gene in a gene-set, and choose the one that is larger in absolute value [8].

### Multiple hypothesis correction

Wherever applicable, we use multiple hypothesis correction based on the Benjamini-Hochberg (BH) approach [51] to obtain corrected *p* values. In case of gene-set analysis, BH approach may result in a conservative estimate of the false discovery rate (FDR) because of overlapping gene-sets that have highly correlated genes. We used the BH method only as a demonstration of statistical power.

## Results

Whole transcriptome expression profile analysis usually focuses on a gene-level analysis by combining gene expression data over all transcripts of a gene. This approach has a significant limitation in that it fails to exploit expression patterns across the transcripts by not explicitly identifying effects that differ among the gene transcripts. Marginal analyses of individual gene transcripts may also lead to a proliferation of hypotheses tested, which can negatively impact the power of biomarker discovery. Popular method used to combine *p* values such as Fisher’s approach assume independence among all the transcripts of a gene, which may not be entirely true in this case. We address the aforementioned issues by proposing two score-test based approaches, one to discover differentially expressed genes and another to identify differentially enriched gene-sets. Score test-based approaches do not require parameter estimation under the alternative hypothesis. As a result, model parameters only have to be estimated once per genome, significantly decreasing computation time. Further, our score-based approaches only require estimation of the first two moments of the random effects, and therefore are robust to misspecification of the random effect distribution [12].

## Evaluating our method to identify differentially expressed (DE) genes using simulated data

We evaluated our method to detect DE genes using two simulation studies. Briefly, each Monte Carlo simulated dataset from the first simulation study was comprised of data for a single gene, whose expression is measured across 5 or 10 transcripts in 50 paired individuals. Each individual pair’s radiation status is either a zero or a one indicating before and after radiotherapy, respectively. Since the transcript-specific effect is modeled as a random effect, a test of whether there is any transcript-specific effect due to radiation is equivalent to testing whether the variance of the random effect (γ) is zero. Thus, our model to detect DE genes involves testing two scalar parameters in β and γ. Simulations under the null hypothesis (no effect of radiation on overall gene expression) confirm that our method has the right type I error. More details in the supplementary section.

Power simulations were performed by varying the following parameters-1) additive effect of radiation (β), 2) the proportion of variation explained by the interaction effect between radiation and transcript (*PV E* γ) and 3) the number of transcripts. The results in table 1 shows that our method significantly outperforms transcript-by-transcript paired t-test and Wilcoxon test (a non-parametric alternative to t-test) in all simulated situations. However, the gene-level paired t-test seems to work the best when there is an overall shift in gene expression due to radiation but absence of any transcript-specific effects.

**Table 1:**
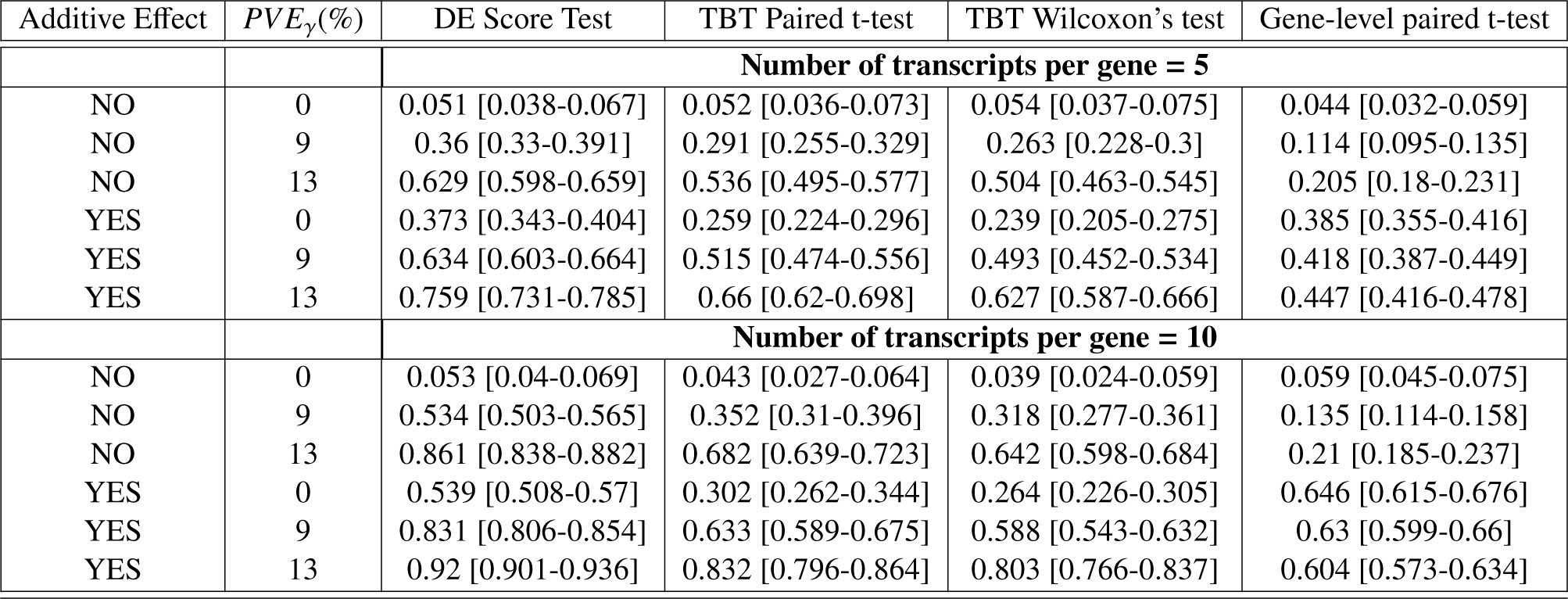
DE of genes - Simulation results at 5% FDR with 95% confidence interval. We varied additive effect i.e. average effect of radiation on the whole transcriptome and proportion of variation explained by γ i.e. radiation × transcripts interaction effect. Our score test is referred to as “DE Score Test”.

| Additive Effect | $PVE_\gamma(\%)$ | DE Score Test | TBT Paired t-test | TBT Wilcoxon's test | Gene-level paired t-test |
| --- | --- | --- | --- | --- | --- |
| **Number of transcripts per gene = 5** |  |  |  |  |  |
| NO | 0 | 0.051 [0.038-0.067] | 0.052 [0.036-0.073] | 0.054 [0.037-0.075] | 0.044 [0.032-0.059] |
| NO | 9 | 0.36 [0.33-0.391] | 0.291 [0.255-0.329] | 0.263 [0.228-0.3] | 0.114 [0.095-0.135] |
| NO | 13 | 0.629 [0.598-0.659] | 0.536 [0.495-0.577] | 0.504 [0.463-0.545] | 0.205 [0.18-0.231] |
| YES | 0 | 0.373 [0.343-0.404] | 0.259 [0.224-0.296] | 0.239 [0.205-0.275] | 0.385 [0.355-0.416] |
| YES | 9 | 0.634 [0.603-0.664] | 0.515 [0.474-0.556] | 0.493 [0.452-0.534] | 0.418 [0.387-0.449] |
| YES | 13 | 0.759 [0.731-0.785] | 0.66 [0.62-0.698] | 0.627 [0.587-0.666] | 0.447 [0.416-0.478] |
| **Number of transcripts per gene = 10** |  |  |  |  |  |
| NO | 0 | 0.053 [0.04-0.069] | 0.043 [0.027-0.064] | 0.039 [0.024-0.059] | 0.059 [0.045-0.075] |
| NO | 9 | 0.534 [0.503-0.565] | 0.352 [0.31-0.396] | 0.318 [0.277-0.361] | 0.135 [0.114-0.158] |
| NO | 13 | 0.861 [0.838-0.882] | 0.682 [0.639-0.723] | 0.642 [0.598-0.684] | 0.21 [0.185-0.237] |
| YES | 0 | 0.539 [0.508-0.57] | 0.302 [0.262-0.344] | 0.264 [0.226-0.305] | 0.646 [0.615-0.676] |
| YES | 9 | 0.831 [0.806-0.854] | 0.633 [0.589-0.675] | 0.588 [0.543-0.632] | 0.63 [0.599-0.66] |
| YES | 13 | 0.92 [0.901-0.936] | 0.832 [0.796-0.864] | 0.803 [0.766-0.837] | 0.604 [0.573-0.634] |

In the second simulation study, each Monte Carlo dataset, comprised of gene expression data for 50 genes over 50 observations, each gene with unequal number of isoforms, was simulated from a multivariate normal distribution with a known variance-covariance matrix. We varied the mean difference in differential gene expression between the two phenotypes (signal-to-noise ratio), and the proportion of differentially expressed gene-isoforms. At the transcript level, we applied paired t-test and a non-parametric alternative in Wilcoxon’s paired t-test and combined the *p* values over all the transcripts of a gene using Fisher’s method. At the gene-level, we combined the gene expression values by computing either the median or Winsorized mean of all the transcripts within a given gene. Paired t-test was run on this gene-level data. We varied the proportion of genes that are differentially expressed and the signal-to-noise ratio. Statistical power and empirical type I error rates were estimated based on a nominal FDR of 5%. Figure 1 displays the performance of all the methods, measured both in terms of statistical power and area under the curve (AUC). AUC for all the methods was estimated using R package ROCR [13]. We see that our method does well compared to the rest of the methods based on AUC plot. Given how the gene expression data were generated, every gene may have a fraction of transcripts differentially expressed. Consequently, any method for identifying DE genes must account for this transcript-specific variability. By combining gene expression values over all the transcripts of a gene (as evidenced by any gene-level methods), we are not able to fully exploit transcript-specific gene expression patterns. This is evident in Figures 1a and 1b, where the gene-level tests perform poorly compared to the transcript-level tests, including our approach.

**Figure 1:**
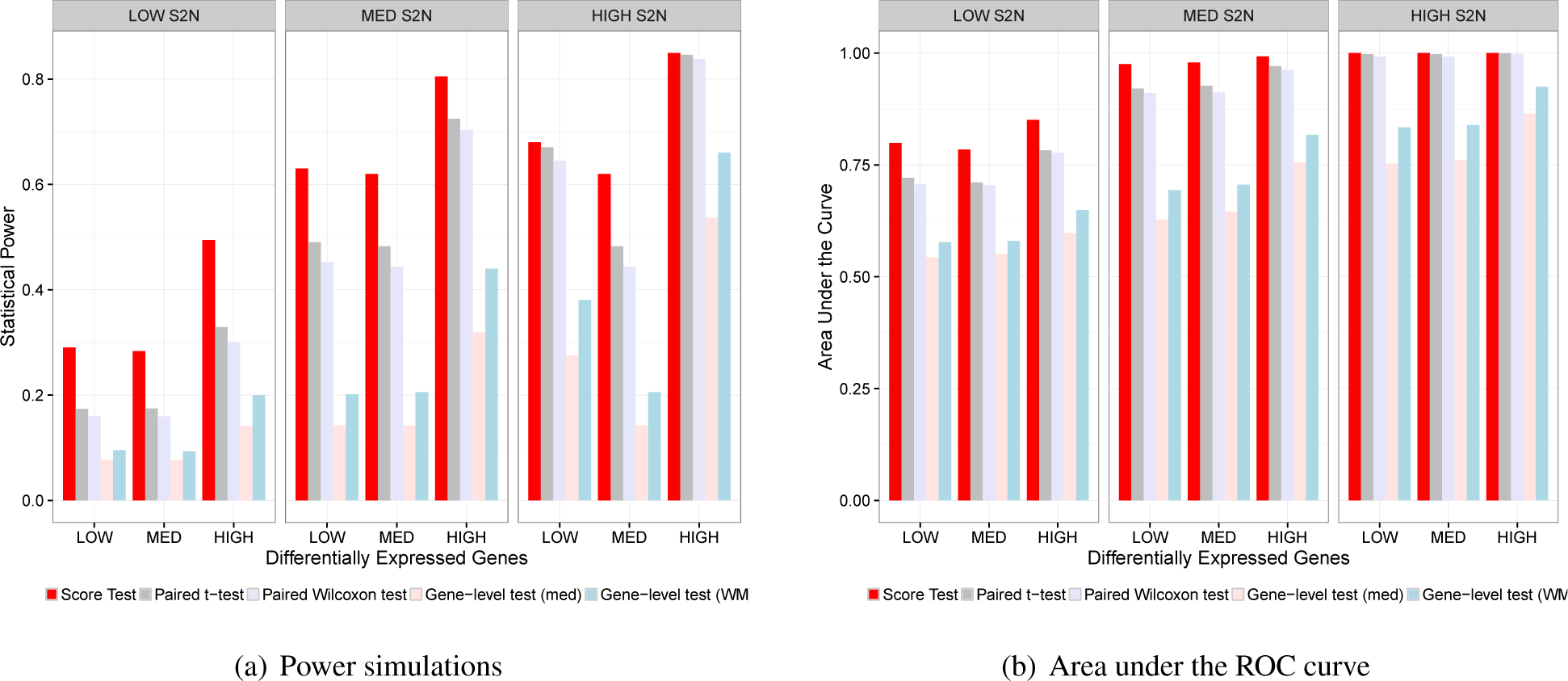
The performance of all the methods in detecting DE genes. A) Bar plot depicting the statistical power of each method under changing number of differentially expressed genes and the mean difference in gene expression (signal-to-noise ratio; S2N) between the two phenotypes (before and after radiation). We compared our method with two transcript-level tests in paired t-test and paired wilcoxon test (*p* values combined at gene-level by Fisher’s method), and with two gene-level tests, where the gene expression values are combined by median and Winsorized mean values followed by a paired t-test. B) Bar plot depicting the area under the curve (AUC) of all the methods under the aforementioned conditions.

## Evaluating our method to identify DE gene-sets using simulated data

We evaluated our method to detect DE gene-sets or pathways using two simulation studies. Briefly, each Monte Carlo simulated dataset from the first simulation study was comprised of data for a single gene-set comprising of 5 genes, whose expression is measured across 3 transcripts in 50 paired individuals. Each individual pair’s radiation status is either a zero or a one indicating before and after radiotherapy, respectively. Since the transcript-specific effect is modeled as a random effect, a test of whether there is any transcript-specific effect on the gene-sets due to radiation is equivalent to testing whether the variances of the random effects (γ and φ) are zero. Thus, our model to detect enriched gene-sets involves testing three scalar parameters in β, γ and φ. Simulations under the null hypothesis (no effect of radiation on overall gene expression) confirm that our method has the right type I error (see supplementary material).

Power simulations were performed by varying the following parameters-1) additive effect of radiation (β), 2) the proportion of variation explained by the interaction effect between radiation and transcript (*PV E* γ) and 3) the proportion of variation explained by the interaction effect between radiation and gene (*PV E* φ). We kept the number of transcripts and genes constant for all these simulations. The results in table 2 show that our method significantly outperforms both transcript-level and gene-level methods. More specifically, our method captures the transcript-specific variability due to radiation within each gene more efficiently than the other tests.

**Table 2:** DE of gene-sets - Gene-set simulation results at 5% FDR with 95% confidence interval. We varied additive effect i.e. average effect of radiation on the whole transcriptome, proportion of variation explained by γ i.e. radiation × transcripts interaction effect, and the proportion of variation explained by φ i.e. radiation × genes interaction effect. Our score test is referred to as “DE Score Test”.

| Additive Effect | $PVE_\gamma(\%)$ | $PVE_\phi(\%)$ | Gene-set Score Test | TBT Paired t-test | TBT Wilcoxon's test | Gene-level paired t-test |
| --- | --- | --- | --- | --- | --- | --- |
| NO | 0 | 0 | 0.048 [ 0.036-0.063 ] | 0.047 [ 0.03-0.07 ] | 0.044 [ 0.027-0.066 ] | 0.042 [ 0.027-0.061 ] |
| NO | 0 | 7 | 0.546 [ 0.515-0.577 ] | 0.234 [ 0.196-0.275 ] | 0.198 [ 0.162-0.237 ] | 0.316 [ 0.279-0.355 ] |
| NO | 0 | 9 | 0.753 [ 0.725-0.779 ] | 0.384 [ 0.339-0.43 ] | 0.313 [ 0.271-0.358 ] | 0.465 [ 0.424-0.506 ] |
| NO | 7 | 0 | 0.408 [ 0.377-0.439 ] | 0.202 [ 0.166-0.242 ] | 0.17 [ 0.137-0.207 ] | 0.12 [ 0.095-0.149 ] |
| NO | 7 | 7 | 0.756 [ 0.728-0.782 ] | 0.413 [ 0.367-0.46 ] | 0.386 [ 0.341-0.432 ] | 0.376 [ 0.337-0.416 ] |
| NO | 6 | 9 | 0.859 [ 0.836-0.88 ] | 0.558 [ 0.511-0.604 ] | 0.515 [ 0.468-0.562 ] | 0.526 [ 0.485-0.567 ] |
| NO | 9 | 0 | 0.584 [ 0.553-0.615 ] | 0.353 [ 0.309-0.399 ] | 0.294 [ 0.253-0.338 ] | 0.178 [ 0.148-0.211 ] |
| NO | 9 | 6 | 0.806 [ 0.78-0.83 ] | 0.546 [ 0.499-0.592 ] | 0.481 [ 0.434-0.528 ] | 0.415 [ 0.375-0.456 ] |
| NO | 8 | 8 | 0.897 [ 0.876-0.915 ] | 0.655 [ 0.61-0.699 ] | 0.601 [ 0.555-0.646 ] | 0.606 [ 0.565-0.646 ] |
| YES | 0 | 0 | 0.716 [ 0.687-0.744 ] | 0.178 [ 0.144-0.216 ] | 0.167 [ 0.134-0.204 ] | 0.289 [ 0.253-0.327 ] |
| YES | 0 | 7 | 0.801 [ 0.775-0.825 ] | 0.386 [ 0.341-0.432 ] | 0.334 [ 0.291-0.379 ] | 0.483 [ 0.442-0.524 ] |
| YES | 0 | 9 | 0.878 [ 0.856-0.898 ] | 0.542 [ 0.495-0.588 ] | 0.483 [ 0.436-0.53 ] | 0.651 [ 0.611-0.69 ] |
| YES | 7 | 0 | 0.738 [ 0.71-0.765 ] | 0.414 [ 0.368-0.461 ] | 0.365 [ 0.321-0.411 ] | 0.334 [ 0.296-0.374 ] |
| YES | 7 | 7 | 0.876 [ 0.854-0.896 ] | 0.588 [ 0.541-0.634 ] | 0.538 [ 0.491-0.584 ] | 0.549 [ 0.508-0.59 ] |
| YES | 6 | 9 | 0.924 [ 0.906-0.94 ] | 0.654 [ 0.609-0.698 ] | 0.607 [ 0.561-0.652 ] | 0.66 [ 0.62-0.698 ] |
| YES | 9 | 0 | 0.763 [ 0.735-0.789 ] | 0.478 [ 0.431-0.525 ] | 0.438 [ 0.392-0.485 ] | 0.349 [ 0.31-0.389 ] |
| YES | 9 | 6 | 0.88 [ 0.858-0.899 ] | 0.654 [ 0.609-0.698 ] | 0.598 [ 0.551-0.643 ] | 0.57 [ 0.529-0.61 ] |
| YES | 8 | 8 | 0.944 [ 0.928-0.957 ] | 0.727 [ 0.684-0.767 ] | 0.682 [ 0.637-0.725 ] | 0.65 [ 0.61-0.689 ] |

In our second simulation study, each Monte Carlo simulation consisted of 100 genes over 5 observations across the two phenotypes. We generated gene expression data using the same approach as described in the previous section. We simulated 10 gene-sets under both scenarios (with non-overlapping and overlapping genes) and compared the performance of our method with the other gene-set enrichment methods at the gene-level. We varied the sizes of gene-sets between 2 and 10 genes. Gene-level analysis is performed by computing the median gene expression values across all the transcripts within a gene followed by an implementation of gene set variational analysis (GSVA), Pathway Level analysis of Gene Expression (PLAGE), single sample GSEA (ssGSEA) and the combined z-score (ZSCORE). We estimated the empirical type I error rate at 5% FDR both in the presence and absence of any gene overlap among the simulated gene-sets. See supplementary methods for more details. In case on no gene overlap, we simulated 10 gene-sets with varying degrees of gene overlap (20%, 50% and 80%), and varying the signal-to-noise ratio between low, medium and high. We compared the performance of all the methods by measuring statistical power and area under the curve in case of gene-sets with no overlapping genes. In the case where gene-sets shared genes, we measured only statistical power.

Figues 2a and 2b show the performance of all the methods when the gene-sets do not share any genes. Even though, this is not a general scenario, our method is competitive with the rest of the methods. In situations where the power of our method is low (relative to the other methods), the accuracy of our method is high given the AUC values. Figure 3 displays the performance of all the methods when the gene-sets have over-lapping genes or shared genes. This is the most common scenario and our method performs well, in terms of statistical power, in almost all cases.

**Figure 2:**
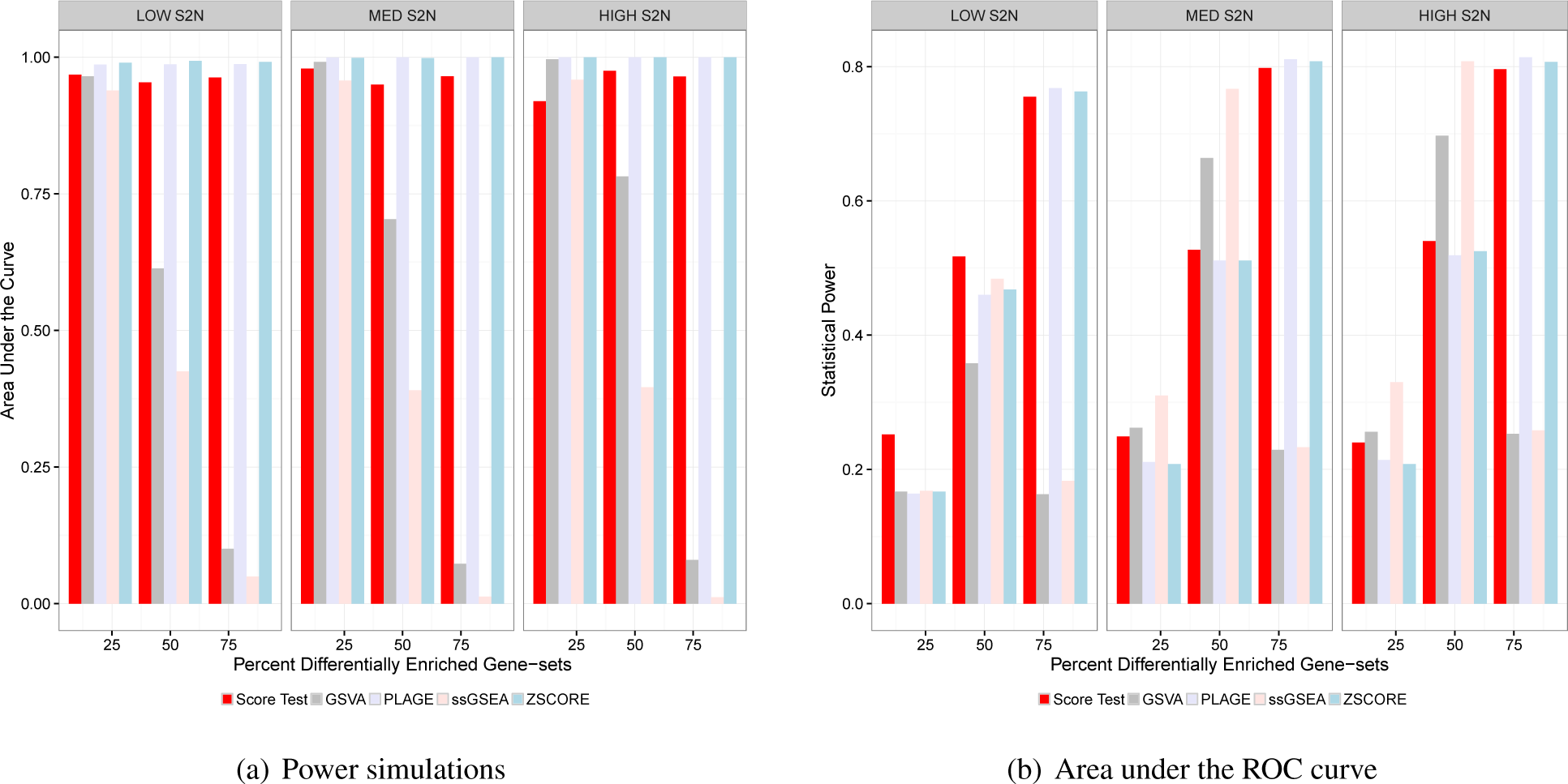
The performance of all the methods in detecting differentially enriched gene-sets when each gene-set is comprised of unique set of genes. A) Bar plot depicting the statistical power of each method under changing number of differentially enriched gene-sets and the mean difference in gene expression (signal-to-noise ratio) between the two phenotypes (before and after radiation). We compared our method with several gene-level tests,by computing the median gene expression values across all the transcripts within a gene. B) Bar plot depicting the area under the curve (AUC) of all the methods under the aforementioned conditions.

**Figure 3:**
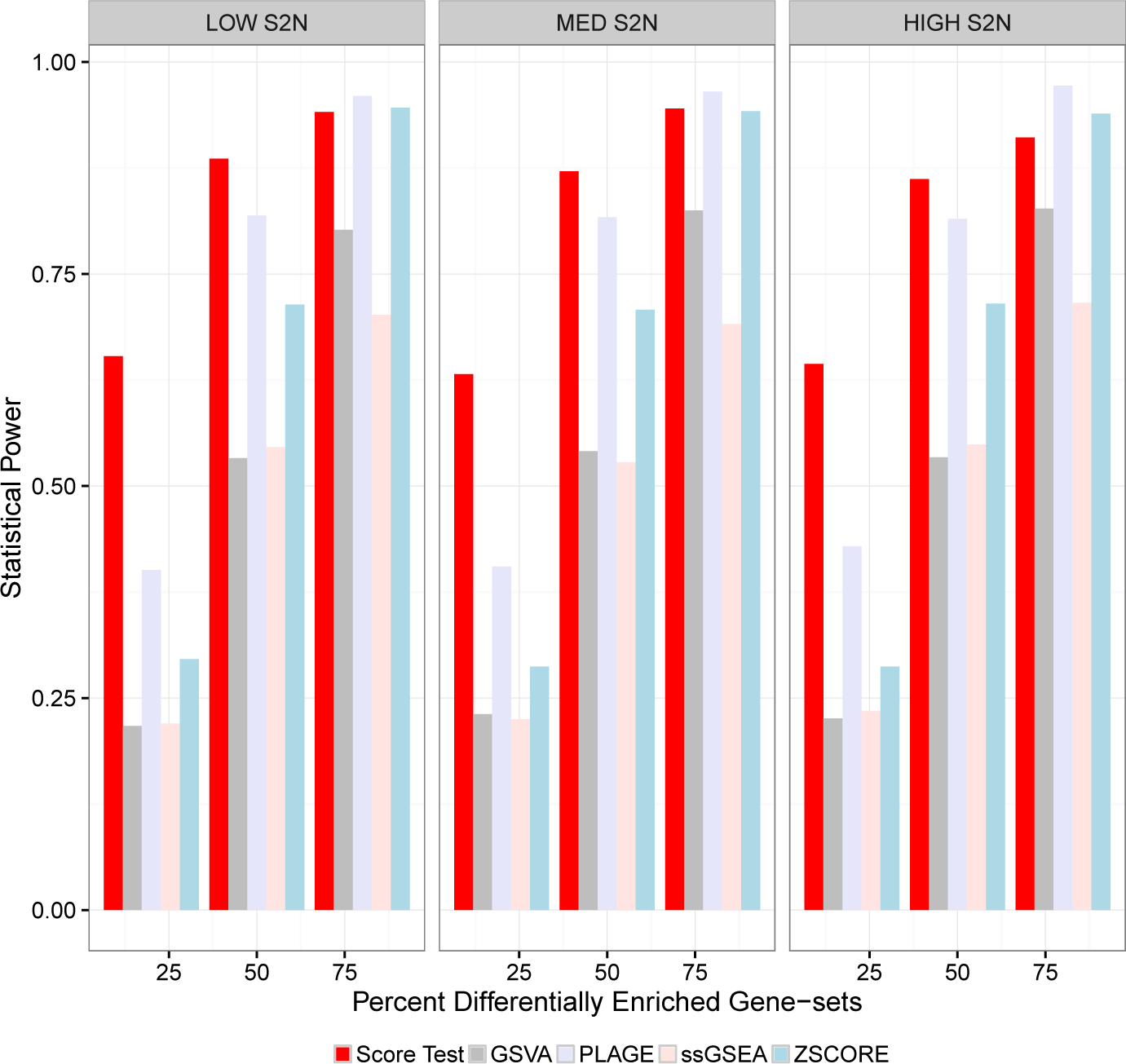
The performance of all the methods in detecting differentially enriched gene-sets when each gene-set is comprised of shared genes. Bar plot depicting the statistical power of each method under changing number of differentially enriched gene-sets and the mean difference in gene expression (signal-to-noise ratio; S2N) between the two phenotypes (before and after radiation). We compared our method with several gene-level tests,by computing the median gene expression values across all the transcripts within a gene.

## Transcriptome-wide response to radiotherapy in breast tumors

### Isoform-specific DE analysis

Transcriptome expression profiling of the early-stage breast cancer patients before and after preoperative radiotherapy using our method has revealed many DE genes. Current methods perform DE analysis at the gene-level and not at the transcript-level. One method performs a standard paired t-test at the transcript-level and combines the resulting *p* values using Fisher’s method [14, 15]. Fisher’s method tests a global null hypothesis that the combined *p* values are jointly significant. However, Fisher’s method assumes that the transcript-level *p* values for each gene are independent. Standard paired t-test followed by Fisher’s method identified 11,944 genes at 5% FDR. Another most commonly used approach is to combine the gene expression values of all transcripts of a gene *a priori* by computing either the median expression values or Winsorized mean expression values (which is robust to any outliers). Paired t-tests were then run on the combined data. These two ways of combining the data identified 4,729 and 3,353 genes, respectively at 5% FDR. Our method identified a total of 12,414 DE genes at 5% FDR, which is more than the ones identified by the aforementioned methods. To assess the biological relevance of the DE genes, we performed a KEGG pathway term enrichment analysis [16] for each set of results separately. KEGG pathways were considered overrepresented if a set of at least three genes from different linked regions is observed to be overrepresented with an adjusted significance level of an adjusted *p* value < 0.05, calculated from a hypergeometric test [17].

The results in table 3 show a list of top 10 signaling pathways that were shown be overrepresented in the dataset without any specifics on the directionality (up- or down-regulation) of the pathway deregulation. For example, PI3K-AKT signaling pathway shown in the table, a potential target for radiosensitizing cancer cells, is one the many pro-survival signaling pathways that get activated by radiation that may lead to suppression of apoptosis, initiation of DNA repair mechanisms and induction of cell-cycle arrest [18]. Together with mTOR signaling pathway, PI3K-AKT are activated in many different cancers. Drugs like rapamycin, CCI-779 and RAD-001 target mTOR signaling pathway while perofisine, PX-866 target AKT pathway [19].

**Table 3:** A list of top 10 over-represented KEGG pathways based on the functional enrichment of our DE gene list.

| KEGG ID | Description | $p$ values | Adjusted $p$ values | $q$ value |
| --- | --- | --- | --- | --- |
| hsa05200 | Pathways in cancer | 4.56E-09 | 1.34E-06 | 7.68E-07 |
| hsa04151 | PI3K-AKT signaling pathway | 3.69E-08 | 5.45E-06 | 3.11E-06 |
| hsa01100 | Metabolic pathways | 1.64E-07 | 1.61E-05 | 9.20E-06 |
| hsa04060 | Cytokine-cytokine receptor interaction | 1.21E-06 | 8.91E-05 | 5.09E-05 |
| hsa04510 | Focal adhesion | 1.59E-06 | 9.38E-05 | 5.36E-05 |
| hsa04630 | JAK-STAT signaling pathway | 3.35E-06 | 0.000164626 | 9.40E-05 |
| hsa04144 | Endocytosis | 9.88E-06 | 0.000386927 | 0.000220904 |
| hsa05166 | HTLV-I infection | 1.13E-05 | 0.000386927 | 0.000220904 |
| hsa04360 | Axon guidance | 1.24E-05 | 0.000386927 | 0.000220904 |
| hsa04210 | Apoptosis | 1.43E-05 | 0.000386927 | 0.000220904 |

### Isoform-specific gene-set analysis

Instead of focusing on individual genes, we turned our focus on functionally related genes referred to as gene-sets or pathways and assess their behavior before and after treatment with radiation. Gene-set enrichment analysis (GSEA) and other similar methods such as Gene Set Analysis (GSA) make use of the entire gene expression profile in order to assess changes of small magnitude in functionally related genes. The aforementioned methods are supervised, which require an *a priori* knowledge of the phenotypes. In contrast, methods such as single sample GSEA [40], GSVA [10], PLAGE [39], and ZSCORE [41] are unsupervised and focus on the relative enrichment of pathways across all the samples rather than the absolute enrichment with respect to a phenotype. All of these methods work at a gene-level and require us to combine gene expression values at the transcript level before any analysis. Our method identified differentially expressed gene-sets by leveraging transcript-specific effects without having to aggregate gene expression over all the probes of a gene. On this basis, we interrogated critical radiation-associated oncogenic signaling pathways and tumor microenvironment signatures and compared the performance of our method with the rest of the methods. Many of the radiation-associated oncogenic signaling pathways were obtained from the hallmark gene-set collection of the Molecular Signatures Database (MSigDB), which were generated by a hybrid approach that combines an automated computational procedure with manual expert curation [20]. All of the investigated 24 oncogenic signaling pathways and 7 tumor microenvironment gene signatures were found to be statistically significant at 5% FDR by our method. All other methods were applied at the gene-level i.e. aggregated gene expression values over all isoforms using median expression values. GSVA identified 22 gene-sets (70.9%), PLAGE identified 26 gene-sets (83.8%), ssGSEA identified 25 gene-sets (80.6%), and ZSCORE identified 27 gene-sets (87%) at 5% FDR. In order to visualize the patterns of pathway regulation, we obtained a matrix containing sample set enrichment scores of all the 31 gene-sets over all the samples using the popular GSA method. From the heat plots in figure 4, radiation induces a hypoxic state, enhances tumor necrosis factors and suppresses angiogenesis. Radiation-induced inflammatory pathways and immune response signatures can be targeted by therapeutics that improve the clinical outcome of radiotherapy by enhancing the radiosensitivity and decreasing any putative metabolic effects.

**Figure 4:**
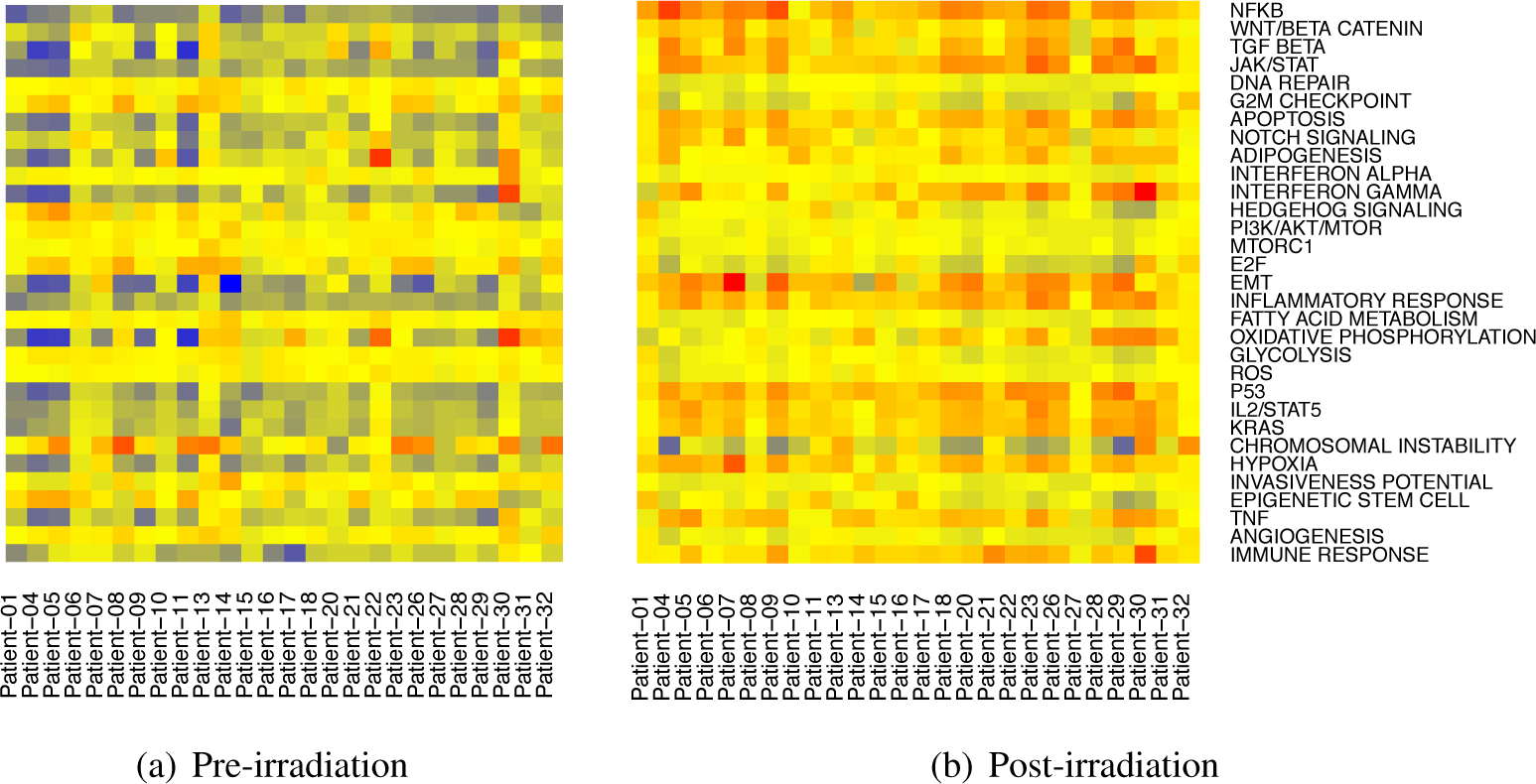
Heat plot showing differentially enriched oncogenic signaling pathways and signatures of tumor microenvironment between patients before and after receiving radiotherapy. The matrix containing sample set enrichment score as computed by the GSA software were used to generate this heat plot. Red indicates a higher collective expression and blue indicates a lower collective expression of genes in that gene-set.

### Discussion

Tailoring a patient’s treatment to exploit an individual’s tumor biology remains an elusive goal in cancer therapy. Similar to cytotoxic therapy, response to radiation in a given population of ‘eligible’ patients is markedly heterogeneous. While chemotherapy serves to address systemic disease, radiation acts as effective local therapy. In many instances, patients resistant to radiation have limited to no options to control local disease [21, 22, 23]; thus, prospectively determining tumor radiosensitivity is important to identify cohorts of patients most likely to respond and to minimize the incidence of radiation-related adverse events in patients who might not otherwise respond. Also, if the molecular underpinnings of radiation response can be elucidated and exploited, the radioresistance of tumors could potentially be abrogated with novel therapeutics. While many mechanisms of radiation resistance, including alterations in DNA repair mechanisms [24], upregulation of pathways regulating angiogenesis [25], apoptosis [26] and cell cycle [27], have been previously described, a comprehensive evaluation of biological events to identify key oncogenic signaling events regulating radiation response, at a genomic and transcriptomic level, is largely unknown. Recent technological advances in quantifying gene expression (i.e. high-throughput sequencing assays) will allow us to interrogate whole exomes or transcriptomes with a higher precision than mRNA expression microarrays thus, overcoming the limitations in detecting and quantifying coding transcript isoforms. However, current statistical methods allow us to interrogate genes and gene-sets at the gene-level by aggregating gene expression across all possible gene isoforms thus, not taking advantage of alternative splicing mechanisms that result in multiple isoforms of the same gene. Overall, our efforts are primarily directed to understanding two very specific aspects - 1) the effect of radiation-induced gene isoform-level variability on gene expression, oncogenic signaling pathways involved in radiation response and tumor microenvironment, and 2) the overall effect of radiation on gene expression. Currently, there are no established methods that leverage gene isoform-specific effects in order to quantify gene expression and investigate tumor biology at a higher resolution. Our methods provide an efficient framework to model transcript-specific and gene-specific effects to map biomarkers association with radiation response. The dataset used here used a high-resolution array-based platform that includes an overwhelming number of gene transcripts in the human transcriptome with >6 million probes targeting coding transcripts, exon-exon splice junctions, and non-coding transcripts. We predict that our methods will also be applicable to gene expression data quantified using RNA-Seq analysis since we make distributional assumptions that preclude their direct application to RNA-Seq count data.

Finally, our methods and analyses are only helpful in generating biological hypotheses, which require substantial verification using *in vitro* and *in vivo* model systems. Eventually, by correctly interpreting these data, we enhance our ability to accurately identify individuals most likely to be resistant to radiotherapy based on the patterns of pathway activation, which further emphasizes the need to identify novel compounds/drugs that could modulate radiation response and function as radiosensitizers.

